# Revising CX3CR1 expression on murine classical and non-classical monocytes

**DOI:** 10.1101/2020.01.28.922534

**Authors:** A. Meghraoui-Kheddar, S. Barthelemy, A. Boissonnas, C. Combadière

## Abstract

In mice, monocytes (Mo) are conventionally described as CX3CR1^low^ classical Mo (CMo) and CX3CR1^high^ non-classical Mo (NCMo) based on the expression level of EGFP in Cx3cr1^*+/EGFP*^ mice and by analogy with human CX3CR1 expression. Although this terminology is widely used, very few works assessed the strict correlation between EGFP reporter and CX3CR1 expression. Using an unsupervised multiparametric analysis of blood Mo in steady state and after sterile peritonitis, we observed no difference in CX3CR1 expression between CMo and NCMo. Our results highlight that despite being a reliable reporter to discriminate Mo subpopulations, EGFP level in Cx3cr1^*+/EGFP*^ mice does not reflect CX3CR1 expression measured by a fluorescently-labeled CX3CL1 chemokine and a CX3CR1 specific antibody. In conclusion, authors should be cautious not to identify murine classical and non-classical Mo as CX3CR1^low^ and CX3CR1^high^ but rather use alternative markers such as the combination of Ly6C and CD43.

## Introduction

Chemokine receptors are key G protein-coupled receptors for immune cell trafficking in inflammation and physiological conditions. They are critical for lymphocytes homing, for normal lymphoid tissue development, for Mo egress from bone marrow (BM) and facilitate organ infiltration of immune cells [1]. Because of their selective expression on leukocyte subsets, they are useful cell surface markers that identify immune cell subtypes like CCR7 for naïve T cells, CXCR5 for T follicular helper lymphocytes, CCR5 and CXCR3 for type 1 lymphocytes or CCR2 and CX3CR1 to discriminate Mo subsets. The latter initially described as an orphan 7-transmembrane domain receptor named alternatively CMKBRL1 [2, 3] or V28 [4] is the specific high-affinity and functional receptor for the chemokine CX3CL1 in mice and human [2, 3, 5, 6]. This chemokine is the sole member of the CX3C chemokine subfamily and was identified in human cells as Fractalkine [7] and in mouse activated brain microglia as neurotactin [8]. It was characterized as a versatile molecule that directed migration of Mo, NK and T cells by its soluble form and regulates adhesion of these cells by its membrane-bound form expressed on endothelial cells [5].

In 2000, Jung and colleagues [9] generated two transgenic strains of mice where the *Cx3cr1* gene was replaced with the gene encoding the enhanced green fluorescent protein (EGFP) commonly referred to as *Cx3cr1*^*+/EGFP*^ and *Cx3cr1*^*EGFP/EGFP*^. This approach allowed the examination of the CX3CR1 expression pattern and migration of cells that normally express this receptor. Based on the green fluorescence and the use of a Fractalkine/NTN-Fc fusion peptide, they confirmed the presence of CX3CR1 on the surface of Mo, part of NK cells, circulating and skin resident DC and microglia. However, the CX3CL1 receptor was absent in resting tissue macrophages (hepatic Kupffer cells, splenic and peritoneal macrophages), astrocytes and oligodendrocytes, neutrophils and eosinophils, B lymphocytes, resting and concanavalin A activated T cells, unlike what has been observed in humans. Nonetheless, recent works clearly demonstrated that terminally differentiated cytotoxic CD8^+^ T cells express CX3CR1 [10]. Consequently to Jung work, *Cx3cr1*^*+/EGFP*^ and *Cx3cr1*^*EGFP/EGFP*^ mice have become widely used and EGFP fluorescence level was used to monitor CX3CR1 expression level in several cell populations and its modulation through time and under several pathological conditions (e.g. inflammation, infection, cancer) [11–15].

In mice, Mo are differentiated in two subsets. It was first achieved on their expression of CCR2, CD62L and CX3CR1 measured by expression of EGFP in cells from *Cx3cr1*^*+/EGFP*^ mice [16]. One Mo subset express CCR2, CD62L and only moderate amounts of EGFP and are known as the ‘inflammatory’ subset, whereas the second that does not express CCR2 or CD62L but display higher expression of EGFP and CD43 is referred as patrolling. In addition, Geissmann *et al.* [17] identified Ly6C as an additional marker of inflammatory Mo in mice. These studies indicated that CCR2^+^CD62L^+^CX3CR1-EGFP^low^Ly6C^+^ mouse Mo correspond to CD14^hi^CD16^−^ classical human Mo, which are also CCR2^+^CX3CR1^low^ and that CCR2^−^CD62L^−^CX3CR1-EGFP^hi^Ly6C^low^ mouse Mo correspond to CD14^low^CD16^+^ human non-classical Mo, which also express large amounts of CX3CR1. These observations were the first to indicate that it would be possible to address the *in vivo* relevance of human Mo heterogeneity by studying mice.

So far, the level of expression of EGFP combined to the detection of Ly6C (or Gr1) marker in *Cx3cr1*^*+/EGFP*^ mice was the most often applied strategy to differentiate CMo, assumed as Ly6C^high^CX3CR1^low^, from NCMo, assumed as Ly6C^low^CX3CR1^high^ [18]. This strategy was and still is commonly used, despite the fact that not all green fluorescent cells in *Cx3cr1*^*+/EGFP*^ mice would be expected to be CX3CR1^+^. Cells that ceased to express the CX3CR1 are likely to harbor residual EGFP because of the extended half-life of the EGFP protein (>24 h) [9]. Green fluorescence in these cells would thus indicate their derivation from CX3CR1-expressing cells but may not reflect the cell expression of the receptor. In fact, Hamon at al. [19] observed in *Cx3cr1*^*+/EGFP*^ mice that the EGFP fluorescent intensity was significantly higher in circulating Ly6C^low^ Mo than Ly6C^high^ Mo. Contrarily to what was observed with EGFP, these cells stained with fluorescently-labelled CX3CL1 showed an equivalent level of CX3CR1. This discrepancy was also observed in the bone marrow. Hence, we addressed the correlation between EGFP and CX3CR1 expression on murine Mo in homeostatic and inflammatory conditions.

## Results and discussion

### CX3CR1 expression does not discriminate Mo subsets

Blood Mo were characterized using an unsupervised analysis based on previously described markers that are preferentially expressed on each subset [16, 17]. The Visualization of t-Distributed Stochastic Neighbour Embedding (viSNE implementation of t-SNE) [21] was used to automatically arrange circulating immune cells according to their expression profile of 16 stained proteins before and after LPS injection, and to visualize all cells in a 2D map where position represents local phenotypic similarity (Fig. 1A). Circulating Mo (purple gate, Fig. 1A) were gated apart from the other circulating immune cells (black cluster, Fig. 1A), on the viSNE map, based on the expression of CD11b, Ly6C, CCR2, CD43, CD36, CD64, NR4A1, and CX3CR1 (Fig. 1B). A second t-SNE map was generated on the former and showed good discrimination of two dominant clusters identified as CMo with preferential expression of Ly6C, CD64, CCR2, CD62L, CD36, CD11b proteins and NCMo with selective expression of CD43 protein and the orphan nuclear receptor Nuclear Receptor Subfamily 4 Group A Member 1 (NR4A1), a transcription factor involved in the differentiation of NCMo from CMo [22], monitored by EGFP reporter from the Nr4A1-EGFP transgenic mouse (Fig. 1C). The unsupervised analysis uncovered that CX3CR1 expression was identical on the two subsets contrariwise to the expected dogma presenting CMo as CX3CR1^low^ and NCMo as CX3CR1^high^ (Fig. 1B, D). Gating on Mo subsets using Ly6C and CD43 markers allowed a better identification of CMo (blue gate and blue cluster, Fig. 1E) and NCMo (green gate and green cluster, Fig. 1E) with a better definition of the intermediate Mo subsets (red gate and red cluster, Fig. 1E).

**Fig. 1:**
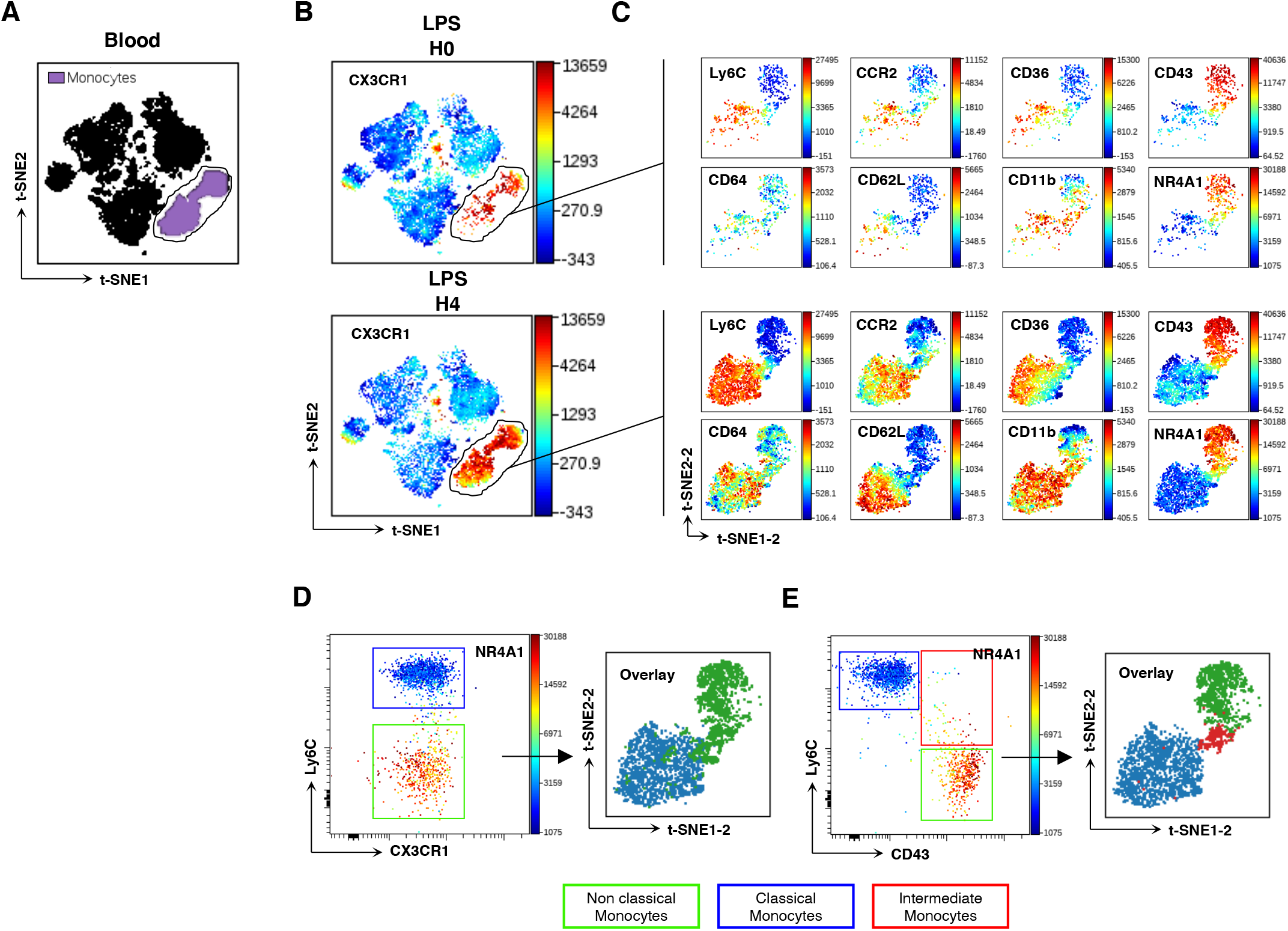
Mo subsets phenotyping. t-SNE was used to automatically arrange circulating immune cells according to their expression profile of 16 stained proteins before (H0) and after (H4) LPS injection in a 2D t-SNE1/t-SNE2 plot **(A)**. The expression of CX3CR1 was presented in a color scale going from blue to red **(B)**. Mo were gated (purple cluster in **A**, black gate in **B)** and visualized using a second axes set t-SNE1-2/t-SNE2-2 **(C)**. The expression of Ly6C, CD64, CCR2, CD36, CD11b, CD43 markers and NR4A1 transcription factor were presented at H0 and H4 after LPS injection in a color scale going from blue to red **(C)**. Whereas gating on Mo subset using CX3CR1-EGFP expression level aloud the discrimination on two subsets **(D)**, the use of CD43 marker in complement to Ly6C permitted the isolation of three subsets expressing different levels of NR4A1 **(E)**. Each subset was back viewed and overlaid on the t-SNE1-2/t-SNE2-2 map **(D, E)**. Dot-plots represent a merged file of three mice per group.

### EGFP fluorescence does not correlate with CX3CR1 expression level on Mo subsets

Because CX3CR1 expression on murine Mo was previously evaluated mainly on EGFP fluorescent reporter of the *Cx3cr1*^*EGFP/+*^ knock-in mice, the expression of the receptor and the fluorescent reporter were considered alike. Here, we investigated whether EGFP expression correlated with the CX3CR1 surface expression, using both its specific ligand and antibody. We first validated a monoclonal anti-CX3CR1-PE (clone: SA011F11) that provide a reliable staining of CX3CR1 in *Cx3cr1*^+/EGFP^ and no staining in *Cx3cr1*^EGFP/EGFP^ mice. CMo (blue gate, Fig. 2A) and NCMo (green gate, Fig. 2A) were conventionally identified based on Ly6C and EGFP expression after the exclusion of neutrophils, T lymphocytes, natural killer cells and eosinophils (CD3^−^SiglecF^−^NK1.1^−^ CD11b^high^Ly6G^−^ cells).

**Fig. 2:**
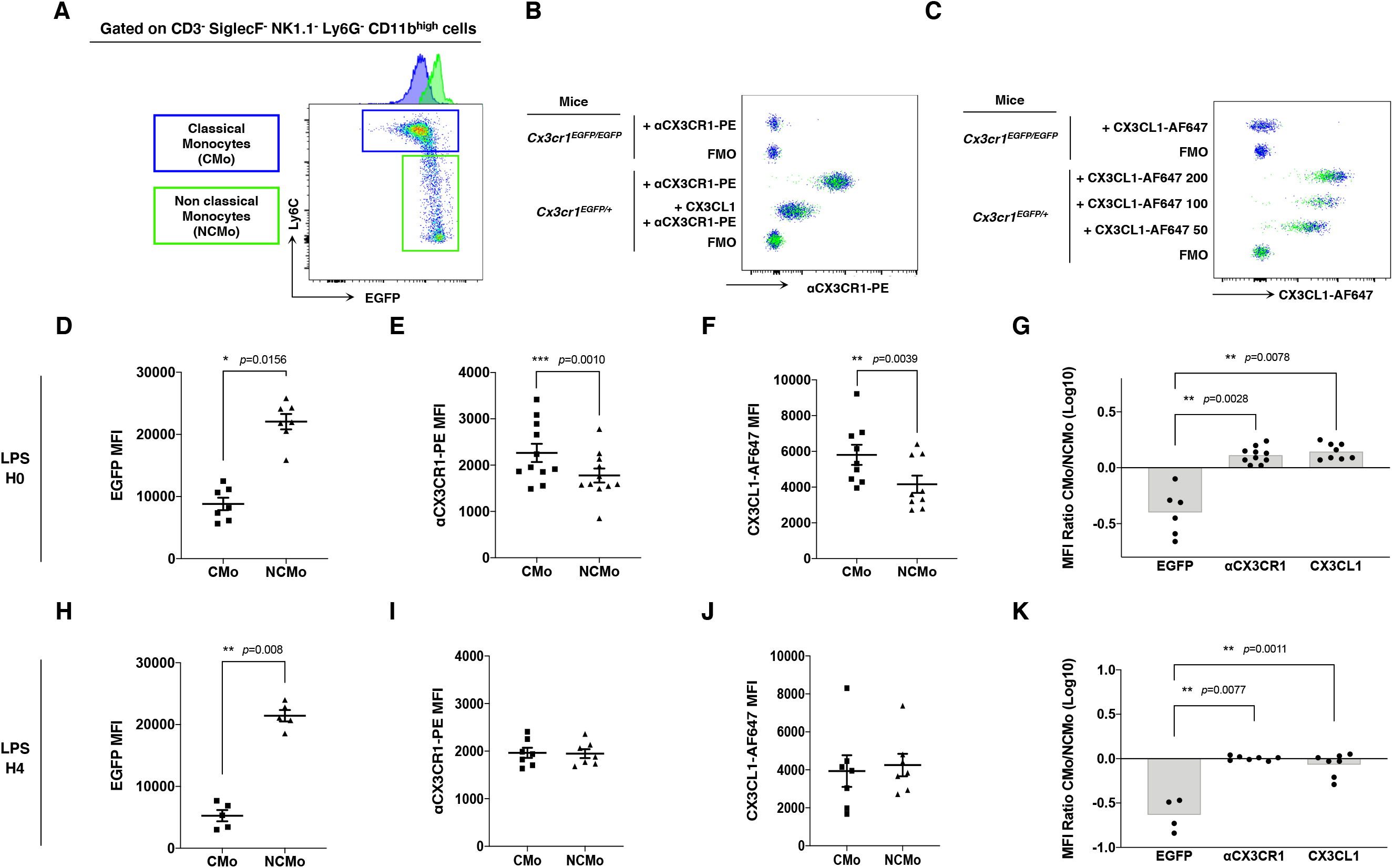
Does EGFP reflects CX3CR1 expression level on monocytes subsets? In *Cx3cr1*^*+/EGFP*^ mice, classical (CMo, blue gate) and non-classical (NCMo, green gate) circulating monocytes were gated on the basis of Ly6C and EGFP expression level as CX3CR1^low^ Ly6C^high^ and CX3CR1^high^ Ly6C^low^ cells respectively **(A)**. CX3CR1 expression level was also assessed using PE conjugated monoclonal antibody specific to CX3CR1 and its specificity was confirmed using a co-incubation with CX3CL1 **(B)**, and by using a chemokine binding assay with 200, 100 and 50nM of CX3CL1-AF647 **(C)**. Staining and binding specificity were assessed using a full minus-one stained sample (FMO) and *Cx3cr1*^*EGFP/EGFP*^ mice. MFI of EGFP **(D**: n=7; **H**: n=5), anti-CX3CR1-PE antibody **(E**: n=11; **I**: n=7) and CX3CL1-AF647 **(F**: n=9; **J**: n=7) were measured on circulating classical (CMo) and non-classical (NCMo) subsets in *Cx3cr1*^*+/EGFP*^ mice at H0 **(D, E, F)** and 4 hours after LPS intra-peritoneal injections **(H, I, J)**. Nonparametric two-tailed Wilcoxon signed-rank test was used to compare differences in MFI from two matched samples. Ratio of CMo MFI over NCMo MFI were calculated at H0 **(G**: n=6-10) and H4 **(K**: n=4-7) after LPS injection.

As historically described, Ly6C^high^ CMo from *Cx3cr1*^*+/EGFP*^ mice, expressed a significantly lower amount of EGFP compared to Ly6C^low^ NCMo (MFI=8805±2652, MFI=22056±3266, respectively, *p*=0.0156) (Fig. 2A, D). However, CMo expressed a higher level of CX3CR1, as it is measured with anti-CX3CR1-PE antibody staining (MFI=2265±648, MFI=1777±500, respectively, *p*=0.0010) (Fig. 2B, E) or with CX3CL1-AF647 staining (MFI=5808±1694, MFI=4164±1447, respectively, *p*=0.0039) (Fig. 2C, F). Based on these results Ly6C^high^ CMo were CX3CR1^+^ and EGFP^low^ whereas Ly6C^low^ NCMo were CX3CR1^+^ and EGFP^high^ with a slightly lower anti-CX3CR1-PE and CX3CL1-A647 MFI in the latter. The ratio of CX3CR1 MFI between CMo and NCMo clearly indicates that the indirect measure of CX3CR1 using the cytosolic EGFP reporter is discordant with the direct measure of CX3CR1 cell surface expression based on both specific chemokine and antibody binding (Fig. 2G). Similar discrepancy was observed on circulating CMo and NCMo 4 hours after intra-peritoneal LPS injection (Fig. 2H-K), indicating that, in both homeostatic and certain inflammatory conditions, CMo and NCMo display different EGFP expression but similar level of CX3CR1 protein expression (Fig. 2K) challenging the original definition of Mo subsets in mice based on CX3CR1 expression.

### CD43 expression identify tissue infiltrating NCMo

We next evaluated whether the combination of CD43 and Ly6C surface markers that allow unambiguous identification of blood CMo from NCMo, would be efficient in others tissues. Mo were harvested from the bone marrow, the spleen and the lungs after blood tissue partitioning by IV injection of a fluorescently labeled anti-CD45 to identify vascular resident cells (CD45+) from tissue-resident cells (CD45-) [23] at homeostasis (Fig. 3A, C) and 4h after LPS intraperitoneal injection (Fig. 3B, D). Similarly to what was observed in the blood, CD43 clearly identify all myeloid cells expressing NRR4A1, in all the tested organs and, in homeostatic and inflammatory conditions (Fig. 3A, B). Blood tissue partitioning revealed that most of the lung NCMo and CMo reside exclusively in the vasculature, with a CD45 MFI equivalent to blood Mo, but in the bone marrow and the spleen, both NCMo and CMo reside within the tissue in steady state as well as 4 hours after LPS inoculation (Fig. 3C, D). NCMo were originally defined as patrolling Mo for their ability to crawl on the luminal side of the endothelium [17]. With these last observations, patrolling denomination should be carefully used and considered as a distinct subset among NCMo that are not exclusively intravascular depending on the tissue.

**Fig. 3:**
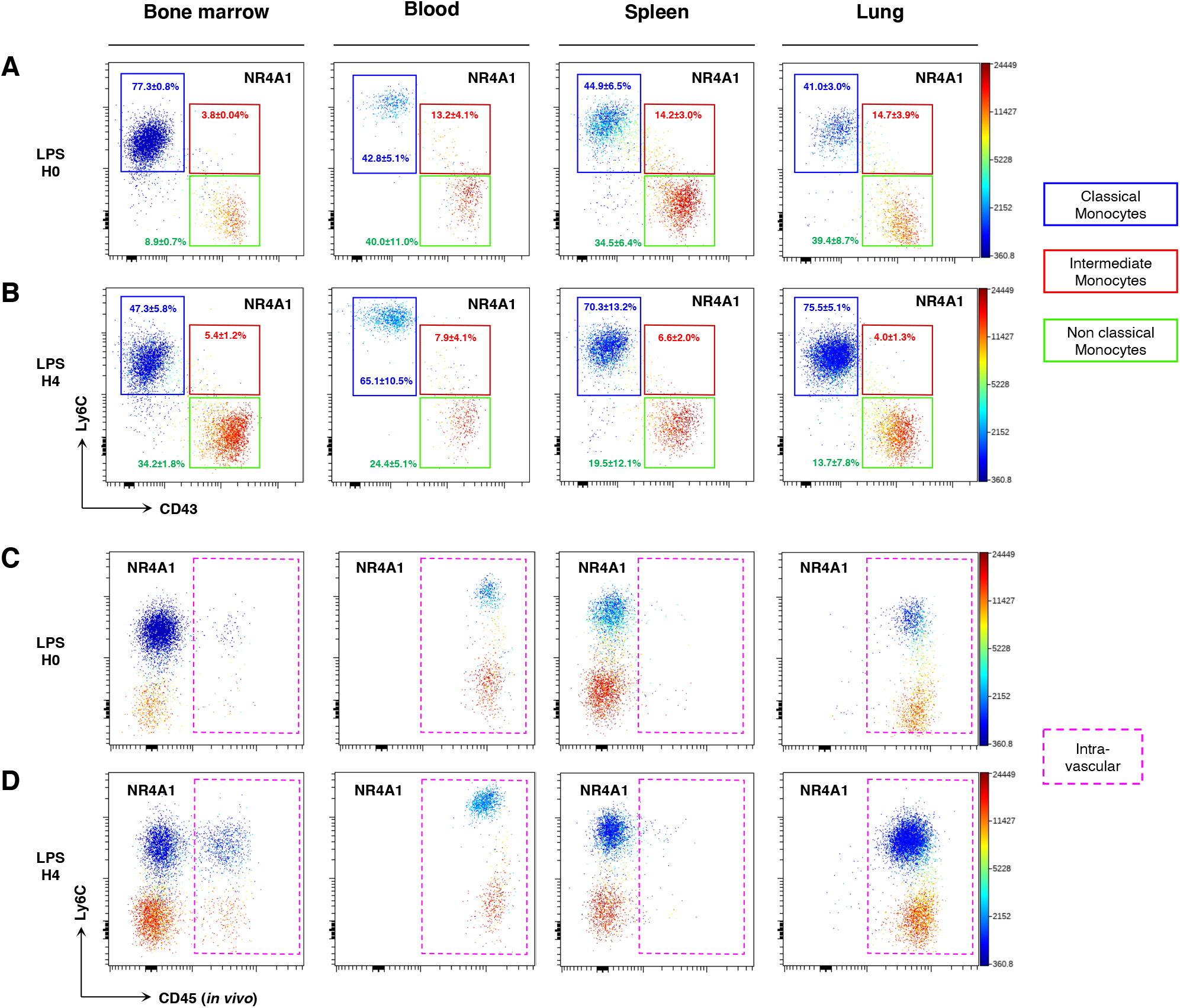
Mo subsets discrimination in tissues. Using Ly6C and CD43 markers, classical (CMo), intermediate (IMo) and non-classial (NCMo) monocyte subsets were gated in bone marrow, blood, spleen and lung before **(A)** and four hours after **(B)** LPS intraperitoneal injection. The distribution of monocytes subsets between the vasculature and the organs parenchyma was discriminated in a Ly6C and CD45 (injected *in vivo*) dot plot before **(C)** and four hours after **(D)** LPS intraperitoneal injection. The level of expression of NR4A1 was presented in each dot plot in a color scale going from blue to red **(A-D)**. Dot-plots are representative of n=3 per group.

In conclusion, we show that Mo EGFP expression level does not correlate with the expression level of CX3CR1 in *Cx3cr1*^*+/EGFP*^ mice, and classical and non-classical Mo definition as CX3CR1^low^ and CX3CR1^high^ respectively should not be used in mice. As a consequence, we propose to refer to an alternative phenotypic strategy using Ly6C and CD43 to identify Mo subsets in the absence of EGFP reporters at homeostasis and inflammation, in blood and tissues.

## Materials and Methods

### Mice

C57Bl6 mice were purchased from Elevage Janvier (Le Genest, Saint Isle, France). Cx3cr1+/EGFP, Cx3cr1EGFP/EGFP [9] and Nr4a1+/EGFP [20] mice were bred in our animal facility “UMS 028 – Phénotypage du petit animal”. All experiments’ protocols were approved by the local ethic committee.

### Sterile Peritonitis models

An LPS-mediated aseptic peritonitis was used in this study. Mice were administered intraperitoneally with 300 ng/kg LPS in phosphate-buffered saline (PBS) or only with PBS for control group.

### Blood tissue partitioning, cells preparation and flow cytometry analysis

Intravascular (IV) CD45 labeling was performed, as previously described [19]. Mice were injected IV with 2 ug of anti-CD45 (clone 30-F11). Two minutes after injection, blood was drawn, mice were sacrificed and needed organs were harvested. Blood was drawn via retro-orbital puncture with heparin and directly stained with antibodies. After staining, erythrocytes were lysed using Pharm Lyse Buffer (BD, Le Pont de Claix, France) following manufacturer instructions. For chemokine binding assays, red blood cells lysing was performed before chemokine binding and antibodies staining. BM cells were harvested by flushing out the thighbone with PBS. Spleen, lung, were harvested and digested in RPMI-1640 medium (Gibco, ThermoFisher, Illkirch, France) containing 1 mg/mL collagenase IV (Sigma Aldrich, Merck, St. Quentin Fallavier) for 30 minutes at 37°C and dissociated through a 30-mm pore cell strainer (Miltenyi Biotec, Bergisch Gladbach, Germany). For chemokine binding assays, cells were incubated with 50, 100 or 200 nM murine CX3CL1-AF647 (Almac, Edinburgh, Scotland) for 30 minutes at 37°C, before antibodies surface staining. CX3CL1-AF647 binding specificity was controlled on Cx3cr1EGFP/EGFP mice. Cell surface staining was performed by incubating for 20 min the freshly prepared cell with Panel-1 or Panel-2 antibodies (Tab. 1) using the adequate dilution for each antibody. The pre-incubated cells with CX3CL1-AF647, were stained with Panel-1 antibodies except anti-CX3CR1 (Tab. 1). Cell suspensions were then washed once using PBS and analyzed directly by flow cytometry. Samples acquisition was performed on the LSRFortessa X-20 Flow cytometry (BD) using FACSDIVA software (BD), and data were analyzed with FlowJo software (BD) and Cytobank analysis platform. (Beckman Coulter, Santa Clara, CA).

**Tab. 1:** Antibodies panels list

| Fluorochrome | Panel-1 |  |  | Panle-2 |  |  |
| --- | --- | --- | --- | --- | --- | --- |
|  | Marker | Clone | Provider | Marker | Clone | Provider |
| PE | CX3CR1 | SA011F11 | Biolegend | CX3CR1 | SA011F11 | Biolegend |
| PECF594 | Ly6G | 1A8 | BD | SiglecF | E50-2440 | BD |
| PECy7 | / | / | / | CD64 | PC61 | Biolegend |
| PerCPCy5.5 | / | / | / | ivCD45 | 30-F11 | BD |
| APC | / | / | / | CD36 | CRF D-2712 | BD |
| APCCy7 | Ly6C | AL-21 | BD | Ly6C | AL-21 | BD |
| BV421 | CCR2 | SA203G11 | Biolegend | CCR2 | SA203G11 | Biolegend |
| BV510 | SiglecF | E50-2440 | BD | MHCII | M5/114 | BD |
| BV605 | / | / | / | CD11c | E50-2440 | BD |
| BV711 | NK1.1 | PK136 | BD | NK1.1 | PK136 | BD |
| BV786 | CD43 | S7 | BD | CD43 | S7 | BD |
| BUV395 | CD11b | M1/70 | BD | CD11b | M1/70 | BD |
| BUV496 | CD3 | 145-2C11 | BD | CD24 | M1/69 | BD |
| BUV563 | / | / | / | Ly6G | 1A8 | BD |
| BUV737 | / | / | / | CD62L | MEL-14 | BD |

### Statistical information

Numeric data are given as mean ± SD with. Nonparametric two-tailed Mann-Whitney test with a significance threshold of alpha (a=0.05) was used to compare differences in mean fluorescence intensity (MFI) between two groups. Nonparametric two-tailed Wilcoxon signed-rank test with a significance threshold of alpha (a=0.05) was used to compare differences in MFI from two matched samples. Statistical tests were performed using commercial statistics software Prism (GraphPad, San Diego, CA).

## Authorship

AMK, AB and CC designed the study. AMK performed experimental work, compiled the data and performed data analysis. SB performed the genotyping of the mice. AB and CC co-authored the manuscript and provided financial support. AMK, AB and CC wrote the manuscript. All authors contributed in reviewing the manuscript.

## Acknowledgments

This work was supported by grants from Inserm, Sorbonne University, Fondation pour la recherche Médicale “Equipe labelisée” and from “Agence Nationale de la Recherche”, project CMOS (CX3CR1 expression on monocytes during sepsis) 2015 (ANR-EMMA-050). AMK was supported by post-doctoral fellowship both from the ANR and FRM.

